# Grb2 binding induces phosphorylation-independent activation of Shp2

**DOI:** 10.1101/859108

**Authors:** Chi-Chuan Lin, Lukasz Wieteska, Kin Man Suen, Arnout Kalverda, Zamal Ahmed, John E. Ladbury

## Abstract

The regulation of phosphatase activity is fundamental to the control of intracellular signalling and in particular the tyrosine kinase-mediated mitogen-activated protein kinase (MAPK) pathway. Shp2 is a ubiquitously expressed protein tyrosine phosphatase and its kinase-induced hyperactivity is associated with many cancer types. In non-stimulated cells we find that binding of the adaptor protein, Grb2, in its monomeric state initiates Shp2 activity independent of phosphatase phosphorylation. Grb2 forms a bidentate interaction with both the N-terminal SH2 and the catalytic domains of Shp2, releasing the phosphatase from its auto-inhibited conformation. Grb2 typically exists as a dimer in the cytoplasm. However, its monomeric state prevails under basal conditions when it is expressed at low concentration, or when it is constitutively phosphorylated on a specific tyrosine residue (Y160). Thus, Grb2 can activate Shp2 and downstream signal transduction, in the absence of extracellular growth factor stimulation or kinase-activating mutations, in response to defined cellular conditions. We identify a polypeptide biotool capable of blocking the Grb2-Shp2 interaction. This peptide down-regulates Shp2 activity *in vitro* and MAPK signalling in a cancer cell line.

## Introduction

The reciprocal process of phosphorylation by kinases, and dephosphorylation by phosphatases, of selected residues regulates the intensity and longevity of intracellular tyrosine kinase-mediated signal transduction. The SH2 domain-containing tyrosine phosphatase 2, Shp2, (aka. PTPN11) plays a prominent role in this process in a multitude of receptor tyrosine kinase (RTK)-mediated signalling pathways, including activation of the extracellular signal-regulated kinase Erk1/2 (aka. mitogen-activated protein kinase, MAPK) pathway (Neel *et al*., 1997; Van Vactor *et al*., 1998; Feng *et al*., 1999; Tonks & Neel 2001; Araki *et al*., 2003). Shp2 is ubiquitously expressed in vertebrate cells and consists largely of, in sequential order: two Src homology 2 (SH2) domains (NSH2 and CSH2 respectively); a protein tyrosine phosphatase (PTP) domain; and a C-terminal tail with two tyrosyl phosphorylation sites (Y542 and Y580) and a proline-rich sequence (residues ^559^PLPPCTPTPP^568^).

Shp2 was the first phosphatase to be identified as a human oncoprotein (Chan & Feng, 2007 and Chan *et al*., 2008), and a large body of experimental and clinical studies has indicated that the hyperactivation of Shp2 contributes to tumour progression in, for example, breast cancer (Bentires-Alj et al., 2004; Bentires-Alj et al., 2006; Bocanegra et al., 2010; Aceto et al., 2012). A great deal of interest has been shown in anti-cancer therapeutic approaches involving down-regulation of Shp2. However, despite extensive investigation, a lack of a mechanistic understanding of its up-regulation in solid cancers has hindered pharmaceutical development. Existing inhibitors targeting phosphatase activity show low selectivity because of the highly conserved amino acid sequences of phosphatase domains.

Crystal structural detail revealed that Shp2 utilises an auto-inhibitory mechanism which prevails under basal conditions (Hof et al., 1998). NSH2 forms an intramolecular interaction with the PTP domain, directly blocking access to the catalytic site, and resulting in a ‘closed’ state. In this state the NSH2 domain adopts a conformation which contorts its phosphopeptide binding cleft. Gain-of-function somatic mutations which result in the abrogation of the interaction between NSH2 and the PTP domain have been shown to be activating (O’Reilly et al., 2000; Chan et al., 2008). Auto-inhibition is released through one of two mechanisms involving the SH2 domains of Shp2. In the first, both N- and CSH2 interact with a binding partner including a phosphorylated bisphosphoryl tyrosine-based activation motif (BTAM) (Lechleider et al., 1993; Sugimoto et al., 1994; Pluskey et al., 1995; Eck et al., 1996; Cunnick et al., 2001). The second, more controversial mechanism occurs under conditions where Shp2 has been phosphorylated, typically by an RTK (Lu et al., 2001; Araki et al., 2003; Keilhack et al., 2005). This induces an intramolecular, bidentate interaction between the two phosphorylated tyrosine residues in the C-terminus of Shp2 and both NSH2 and CSH2 (Lu et al., 2001; Neel et al., 2003; Sun et al., 2013).

We have previously investigated the constitutive control that the adaptor protein growth factor receptor-bound protein 2, Grb2, exerts over Shp2 in non-stimulated cells (Ahmed et al., 2010; Ahmed et al., 2013). Grb2 consists of an SH2 domain sandwiched between two SH3 domains and is integral to several RTK-mediated signalling pathways. Non-phosphorylated Grb2 exists in a concentration-dependent dimer-monomer equilibrium (K_d_ = 0.8 µM; Ahmed et al., 2015). In addition to the result of depletion of cellular concentration (Timsah et al, 2014), monomeric Grb2 (mGrb2) will also prevail when it is phosphorylated on tyrosine 160 (Y160) in the dimer interface (Ahmed et al., 2015). In the absence of growth factor stimulation Grb2 cycles between the phosphorylated mGrb2, and the non-phosphorylated, typically dimeric state. The former is dependent on constitutive, background RTK activity, e.g. fibroblast growth factor receptor 2 (FGFR2; Lin et al., 2012), whilst the latter results from concomitant Shp2 activity (Ahmed et al., 2015).

In this work we provide molecular mechanistic detail on the activation of Shp2 in the absence of the two phosphorylation-dependent mechanisms highlighted above. We show that in non-stimulated cells mGrb2 is able to greatly enhance Shp2 phosphatase activity via a bidentate interaction involving to two discrete interfaces; 1) between the NSH2 domain of Shp2 and the SH2 of Grb2, and 2) between the PTP domain of Shp2 and the CSH3 of Grb2. The binding of mGrb2 releases Shp2 from its auto-inhibited state and results in an increase in the phosphatase activity independent of kinase-induced stimulation. To endorse our mechanistic model we were able to down-regulate Shp2 activity both *in vitro* and in a triple negative breast cancer cell line using a polypeptide that blocks the Grb2 SH2 binding region on the Shp2 NSH2 domain.

## Results

### Shp2 interacts with monomeric Grb2 in the absence of growth factor

Mutation of Y160 to glutamate (a phosphotyrosine charge mimetic) in the Grb2 dimer interface abrogates self-association of the adaptor protein (Ahmed et al., 2015). To characterise the interaction between full-length Shp2 (Shp2_FL_) and monomeric Grb2 (including Y160E mutation, Grb2_Y160E_; Ahmed et al., 2015) we initially used microscale thermophoresis (MST) to measure the affinity of the interaction of full length proteins (K_D_ = 0.33 ± 0.04 µM; Fig 1A). We demonstrated that Grb2_Y160E_ can bind at two discrete sites on Shp2 using bio-layer interferometry (BLI) on the following four GST-tagged phosphatase constructs: Shp2_FL_, the tandem SH2 domains (residues 1-220: Shp2_2SH2_), the PTP domain (221-524: Shp2_PTP_) and a peptide corresponding to the C-terminal 69 amino acids (525-593: Shp2_C69_). Both Shp2_FL_ as well as Shp2_2SH2_ polypeptides were able to interact with Grb2_Y160E_. The truncated Shp2_PTP_ also bound, whilst the C-terminal tail failed to interact (Fig 1B). These two binding sites were confirmed in an *in vitro* pull down experiment in which Grb2_Y160E_ was precipitated by both GST-Shp2_2SH2_ and GST-Shp2_PTP_ (Fig 1C). The interaction with GST-Shp2_PTP_ was less pronounced. Interaction between Grb2_WT_ and the Shp2 constructs appears to be negligible suggesting that, under the experimental conditions, the prevailing dimeric Grb2 is unable to interact. The limited complex formation seen with an extended exposure of the blot is again presumed to be with the low population of monomeric protein at equilibrium (Fig 1C inset).

**Figure 1.**
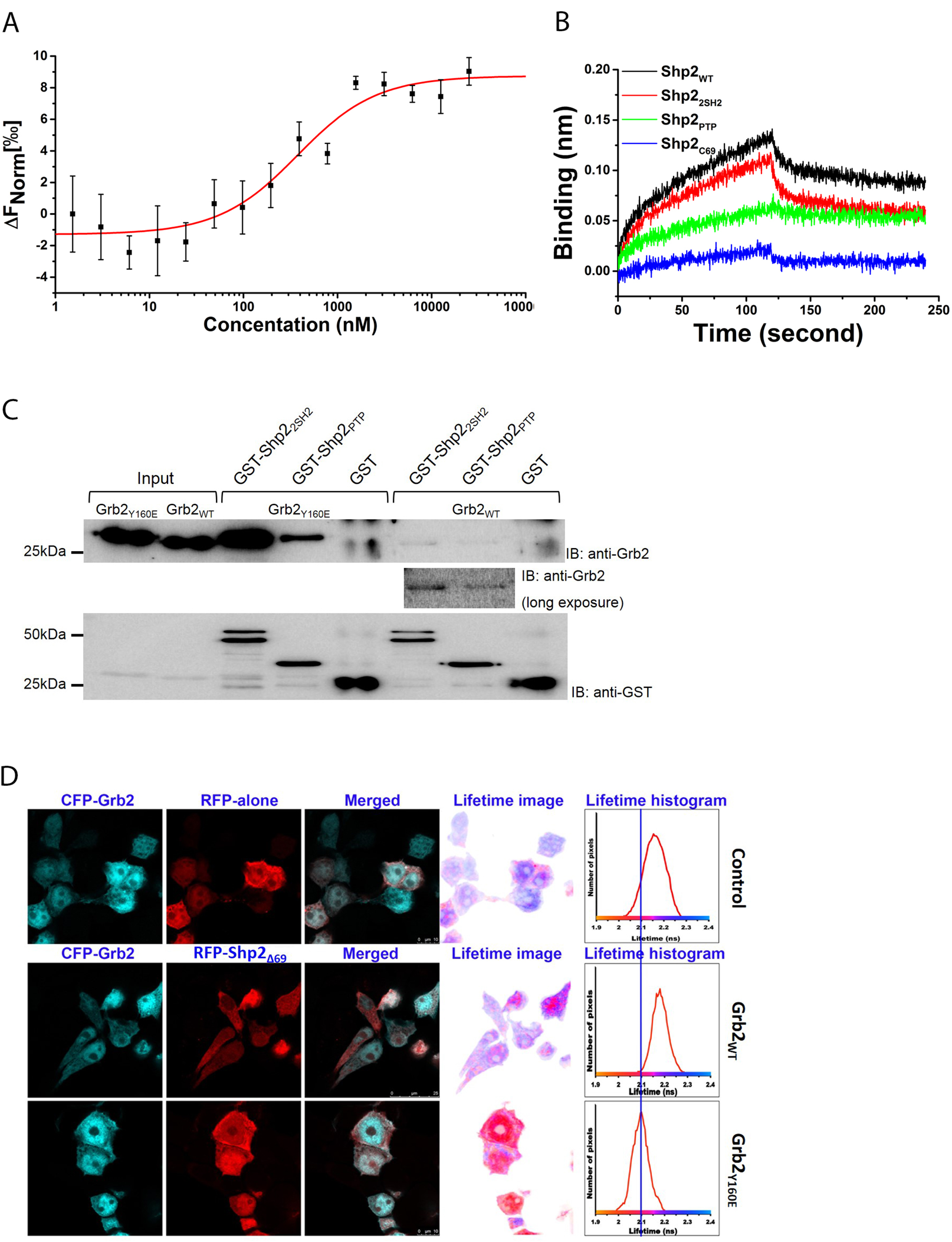
Shp2 interacts with monomeric Grb2 in a phosphorylation-independent manner. A MST measurement of full length Shp2 binding to fluorescent labelled monomeric Grb2. For further details, see Table 1 and Materials and Methods. B BLI characterisation of individual Shp2 domains binding to GST-Grb2_Y160E_ immobilised on a GST sensor. GST-Grb2_Y160E_ was captured and the sensor and 10µM of each Shp2 protein was used to test the interaction. Black: Shp2_WT_, Red: Shp2_2SH2_, Green: Shp2_PTP_, Blue: Shp2_C69_. The BLI sensorgram indicates that the Shp2_2SH2_ and Shp2_PTP_ mediate the interaction with Grb2. C GST pulldown experiment using GST tagged Shp2_2SH2_ and GST tagged Shp2_PTP_ to precipitate monomeric or dimeric Grb2 (Grb2_Y160E_ or Grb2_WT_ respectively). The pulldown results clearly demonstrate the interaction of Shp2 with monomeric Grb2. D Fluorescence lifetime imaging microscopy (FLIM) displaying fluorescence resonance energy transfer (FRET) between CFP-Grb2_WT_ and RFP alone control (top); CFP-Grb2_WT_ and RFP-Shp2_Δ69_ (middle); Grb2_Y160E_ and RFP-Shp2_Δ69_ (bottom). The lifetime-image was generated using false colour range pixel-by-pixel lifetime value corresponding to the average lifetime shown in the histograms.

**Table 1:** Binding affinities (Kd, µM) obtained from this study using MST. Target proteins were fluorescently labelled. Serial dilutions of ligands were titrated in order to determine the binding affinity.

| Target protein | Ligand | $K_d$ ($\mu\text{M}$) |
| --- | --- | --- |
| Grb2<sub>Y160E</sub> | Shp2<sub>FL</sub> | $0.33 \pm 0.04$ |
| Grb2<sub>Y160E</sub> | Shp2<sub>2SH2</sub> | $0.28 \pm 0.03$ |
| Grb2<sub>R86A/Y160E</sub> | Shp2<sub>2SH2</sub> R32A/R138A | $0.15 \pm 0.01$ |
| Shp2<sub>NSH2</sub> | Grb2<sub>SH2</sub> | $33.8 \pm 4.5$ |
| Shp2<sub>NSH2</sub> | Grb2<sub>NSH3</sub> | $422 \pm 21.5$ |
| Shp2<sub>NSH2</sub> | Grb2<sub>CSH3</sub> | $207 \pm 16.8$ |
| Shp2<sub>PTP</sub> | Grb2<sub>Y160E</sub> | $0.30 \pm 0.03$ |
| Grb2<sub>CSH3</sub> | VLHDGD<sup>297</sup>PNEP<sup>300</sup>VSDYIN | N.B. |
| Grb2<sub>CSH3</sub> | TKCNNS<sup>322</sup>KPKK<sup>325</sup>SYIATQ | $173 \pm 6.89$ |
| Grb2<sub>CSH3</sub> | VERGKS<sup>366</sup>KCVK<sup>369</sup>YWPDEY | $122 \pm 7.24$ |
| Grb2<sub>CSH3</sub> | YGVMRV<sup>386</sup>RNVK<sup>389</sup>ESAAHD | N.B. |
| Grb2<sub>CSH3</sub> | AHDYTL<sup>399</sup>RELKLSK<sup>405</sup>VGQ | N.B. |
| Grb2<sub>CSH3</sub> | WPDHGV<sup>429</sup>PSDP<sup>432</sup>GGVLDF | $18.1 \pm 0.188$ |
| Shp2<sub>NSH2</sub> | polypeptide inhibitor (PI) | $7.24 \pm 0.52$ |
| Shp2<sub>PTP</sub> | PI | N.B. |
| Grb2<sub>Y160E</sub> | PI | N.B. |
| Shp2<sub>NSH2</sub>+PI | Grb2<sub>Y160E</sub> | 101 ± 6 |
| Shp2<sub>NSH2</sub> | Grb2<sub>Y160E</sub> | 2.3 ± 0.2 |
| Shp2<sub>NSH2</sub>+PI | Shp2<sub>PTP</sub> | 0.25 |
| Shp2<sub>NSH2</sub> | Shp2<sub>PTP</sub> | 0.55 |

To measure intracellular binding of Shp2_Δ69_ (deleted of residues 525-593) to Grb2_Y160E_ we used fluorescence lifetime imaging microscopy (FLIM) to detect stable complexes through fluorescence resonance energy transfer (FRET) between fluorophore-tagged proteins transfected into HEK293T cells under serum-starved conditions. FRET was confirmed by the left-shift of the fluorescence lifetime from the control lifetime measurements of donor (CFP-Grb2) in the presence of non-specific RFP-alone acceptor (2.2 nsec; (Fig 1D). No interaction between CFP-Grb2_WT_ and RFP-Shp2_Δ69_ was observed since the fluorescent lifetime remains largely unaffected. Grb2_WT_ exists in a monomer-dimer equilibrium and CFP-Grb2_WT_ overexpression would favour dimer at the elevated adaptor protein concentration, hence limiting the availability of mGrb2 to interact with Shp2_Δ69_. To circumvent concentration-dependent dimerization, we used the monomeric CFP-Grb2_Y160E_ in the FLIM binding assay. FRET between CFP-Grb2_Y160E_ and Shp2_Δ69_ shows a left-shifted fluorescence lifetime by 100 psec shorter than the RFP-alone control or CFP-Grb2_WT_ (Fig 1D lower panel). This clearly demonstrates that only the monomeric Grb2_Y160E_ interacts with Shp2. Immunoprecipitation further revealed that Grb2_Y160E_ constitutively associates with Shp2_FL_ in the absence of ligand stimulation (Fig EV1).

### The SH2 domain of Grb2 binds to the N-terminal SH2 domain of Shp2

The affinity of Shp2_2SH2_ binding to monomeric Grb2_Y160E_ was determined by MST (K_d_ = 0.28 ± 0.03 µM; Fig 2A). The binding of the pY-mimetic glutamate of Grb2_Y160E_ to either of the SH2 domains of Shp2_2SH2_ was ruled out because on mutating arginine residues normally required for pY binding in the respective SH2 domains of both proteins (R32A/R138A; Shp2_2SH2 R32A/R138A_ and R86A; Grb2_R86A/Y160E_), Grb2_R86A/Y160E_ was still capable of binding to Shp2_2SH2 R32A/R138A_ with similar affinity to the wild type Shp2_2SH2_ construct (K_d_ = 0.15 ± 0.01 µM; Fig 2B). Having observed the Shp2_2SH2_ interaction with Grb2, we sought to identify whether an individual domain was responsible for the critical contact. GST-tagged Shp2_2SH2 R32A/R138A_ as well as the individual SH2 domains (N-terminal SH2 domain: Shp2_NSH2_ and C-terminal SH2 domain: Shp2_CSH2_) were used in pull down experiments in which the NSH2, and not the CSH2, domain of Shp2 was shown to be sufficient for Grb2 binding (Fig 2C). Re-probing the blot with an anti-pY antibody revealed that the interaction was not mediated through phosphorylated tyrosine(s) or glutamate on Grb2_Y160E_. This was further confirmed through *in vitro* binding assays using two phosphopeptides corresponding to the phosphorylatable tyrosine residues on Grb2 (pY160 and pY209) which show negligible interaction with Shp2_2SH2_. (Fig EV2A). Our data therefore are consistent with a non-canonical, phosphorylation-independent interaction between Shp2_2SH2_ and mGrb2.

**Figure 2.**
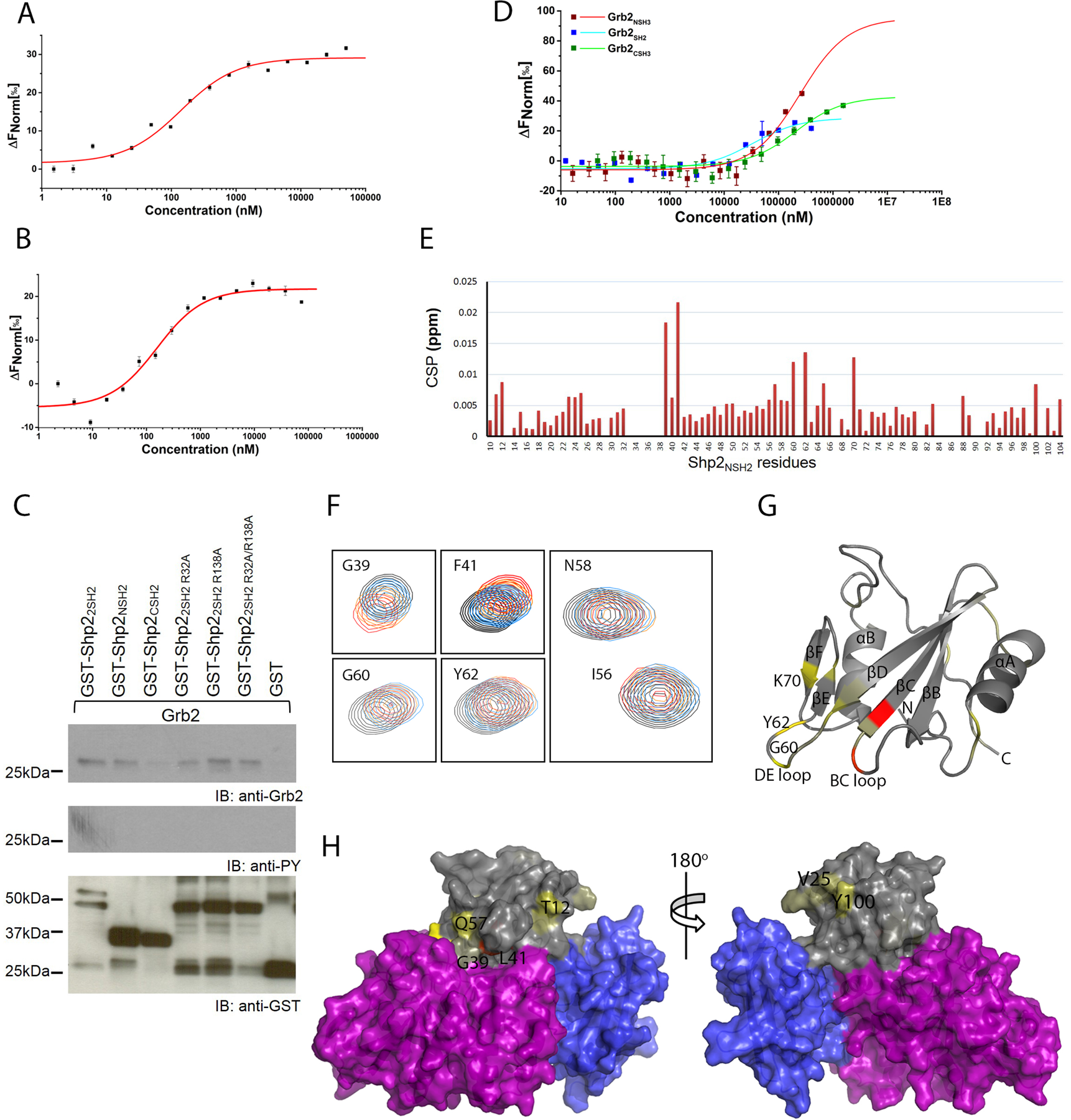
Identification of binding domain on Grb2 and Shp2_2SH2_ domains. A MST measurement of Shp2_2SH2_ binding to fluorescent labelled Grb2_Y160E_. For further details, see Table 1 and Materials and Methods. B MST measurement of Shp2_2SH2 R32A/R138A_ binding to fluorescent labelled Grb2_R86A/Y160E_. This experiment confirms a phosphorylation-independent interaction. C GST pulldown experiment using GST tagged Shp2_NSH2_, Shp2_CSH2_, Shp2_2SH2_ and its SH2 domain deficient mutants. The pulldown results indicate that SH2 domain deficient mutants have no effect on precipitating Grb2 and identify that the NSH2 of Shp2 mediates the interaction with Grb2. Together with the negative result from the phosphotyrosine immunoblotting of precipitated Grb2, it is further confirmed that the Shp2_2SH2_ – Grb2 interaction is phosphorylation-independent. D MST characterisation of Grb2 individual domains binding to fluorescent labelled Shp2_NSH2_. The result indicate that the SH2 domain of Grb2 predominately binding to Shp2_NSH2_. E Interaction of ^15^N-labelled Shp2_NSH2_ and Grb2_SH2_ determined by NMR chemical shift perturbation (CSP) plot for the interaction of Grb2_SH2_ to ^15^N-labelled Shp2_NSH2_. Minimal chemical shift perturbation upon Grb2_SH2_ mapped on Shp2_NSH2_ sequence. F Comparison of Shp2_NSH2_ chemical shifts in the presence and absence of Grb2_SH2_. Overlay of a region of the ^15^N-^1^H HSQC (heteronuclear single-quantum coherence) spectra of Shp2_NSH2_ and Grb2_SH2_ complex (red) and of Shp2_NSH2_ alone (black). G Mapping on the Shp2 N-SH2 domain of the (PDB: 2SHP) of the consensus residues (painted light yellow to red) exhibiting strong CSPs upon Grb2_SH2_ binding. H Mapping on the surface of Shp2 (PDB: 2SHP) of the consensus residues (painted light yellow to red) exhibiting strong CSPs upon Grb2_SH2_ binding. The Shp2_NSH2_ domain is in grey, Shp2_CSH2_ domain is in blue, and the SHp2_PTP_ domain is in purple. This identify the potential binding sites on Shp2_NSH2_ for Grb2_SH2_ binding.

Having established the role of Shp2_NSH2_ in binding to unphosphorylated Grb2_Y160E_, we attempted to establish which domain(s) of Grb2 was required for recognition of the phosphatase. GST-tagged Shp2_NSH2_ was captured on a GST BLI sensor and was exposed to the isolated SH2, NSH3 and CSH3 domains of Grb2 (Grb2_SH2_, Grb2_NSH3_ and Grb2_CSH3_ respectively). The data indicate that Grb2_SH2_ binds to Shp2_NSH2_ (Fig EV2B), accompanied by a weaker interaction with Grb2_CSH3_. MST experiments on the same isolated domains of Grb2 binding to Shp2_NSH2_ confirmed that the interaction with Grb2_SH2_ was dominant (K_d_ = 33.8 ± 4.5 µM). Interactions with both the SH3 domains of Grb2 were substantially weaker (Grb2_NSH3_^;^ K_d_ = 422 ± 21.5 µM, and Grb2_CSH3_; K_d_ = 207 ± 16.8 µM (Fig 2D).

To assess the impact of the Grb2_SH2_ domain binding to Shp2 NSH2 domain, we carried out NMR titration experiments based on 2D (^1^H, ^15^N) HSQC spectra of ^15^N-Shp2 NSH2 domain in the absence, and the presence of increasing concentration of unlabelled Grb2_SH2_ domain. Peak assignments of Shp2_NSH2_ backbone ^1^H-^15^N resonances were derived (81% assignment coverage; Fig EV3). Addition of Grb2_SH2_ led to chemical shift perturbations (CSPs) for several resonances, indicating disruption of local chemical environments of specific amino acids as a result of complex formation (Fig 2E). Comparison of the spectra of the free Shp2_NSH2_, and the Shp2_NSH2_-Grb2_SH2_ complex showed that the average CSPs are relatively small. However, a limited number of residues show pronounced changes (Fig 2F). The CSPs (>0.0075 ppm) were mapped onto a crystal structural representation of Shp2_NSH2_ (PDB code: 2SHP; Figs 2G and 2H). From this it is possible to interpret a mechanism for Grb2_SH2_ domain binding. The reported crystal structural data of Shp2 (PDB: 2SHP) reveal a non-phosphorylated auto-inhibited structure in which NSH2 directly interacts with the PTP domain, blocking the access of phosphorylated substrates. We see that a number of residues which are not within the Shp2 NSH2-PTP interface are perturbed by binding Grb2_SH2_ (T12, V25, Q57 and Y100; Fig 2H) and are likely to represent a discrete and extended interface formed between Grb2 and Shp2. In addition to this we observe that a subset of residues, that are within the intramolecular auto-inhibited interface also show CSPs (G39, F41, G60, Y62 and K70; Figs 2F, and 2G). For instance, G60 and Y62 are located on the DE loop, which inserts into the PTP domain catalytic site interacting with residues of the catalytic loop of Shp2 (Hof et al., 1998). This suggests that binding of Grb2_SH2_ is also able to induce a distal conformational change in Shp2_NSH2_ capable of disrupting the auto-inhibited intramolecular interface releasing the ‘closed’ structure and hence activating the phosphatase.

### The CSH3 domain of Grb2 interacts with Shp2_PTP_

Data shown in Fig 1B and 1C revealed that, as well as binding to Shp2_2SH2_, Grb2_Y160E_ also formed an interaction with Shp2_PTP_. We explored this further using MST and found that the isolated Shp2_PTP_ binds Grb2_Y160E_ with a similar affinity to the interaction with Shp2_2SH2_ (K_d_ = 0.30 ± 0.03 µM; Fig 3A). To identify which domain of Grb2 interacts with Shp2_PTP_ we used GST-fused Grb2_SH2_, Grb2_NSH3_ and Grb2_CSH3_ constructs to precipitate recombinant Shp2_PTP_. The pulldown indicated that only Grb2_CSH3_ can bind to the PTP domain (Fig 3B). SH3 domains recognise sequences usually incorporating the motif PXXP (where X is any amino acid), although the atypical R/KXXK motif has also been associated with SH3 domain recognition. We found that there are two PXXP motifs (^297^PNEP^300^ and ^429^PSDP^432^) and four R/KXXK SH3 domain-binding motifs (^322^KPKK^325^, ^366^KCVK^369^, ^386^RNVK^389^ and ^399^RELKLSK^405^) within the PTP domain which could potentially serve as Grb2_CSH3_ binding sites. We therefore performed MST assays using Grb2_Y160E_ and six 16-residue peptides incorporating these sequences. Binding data showed that the Grb2_CSH3_ had the highest affinity for the ^429^PSDP^432^ on the Shp2_PTP_ (K_d_ ∼ 18 µM; Fig 3C and Table). This interaction is approximately 60-fold weaker than the interaction between the intact PTP domain and Grb2_Y160E_ suggesting that the peptides do not entirely represent the full site of interaction.

**Figure 3.**
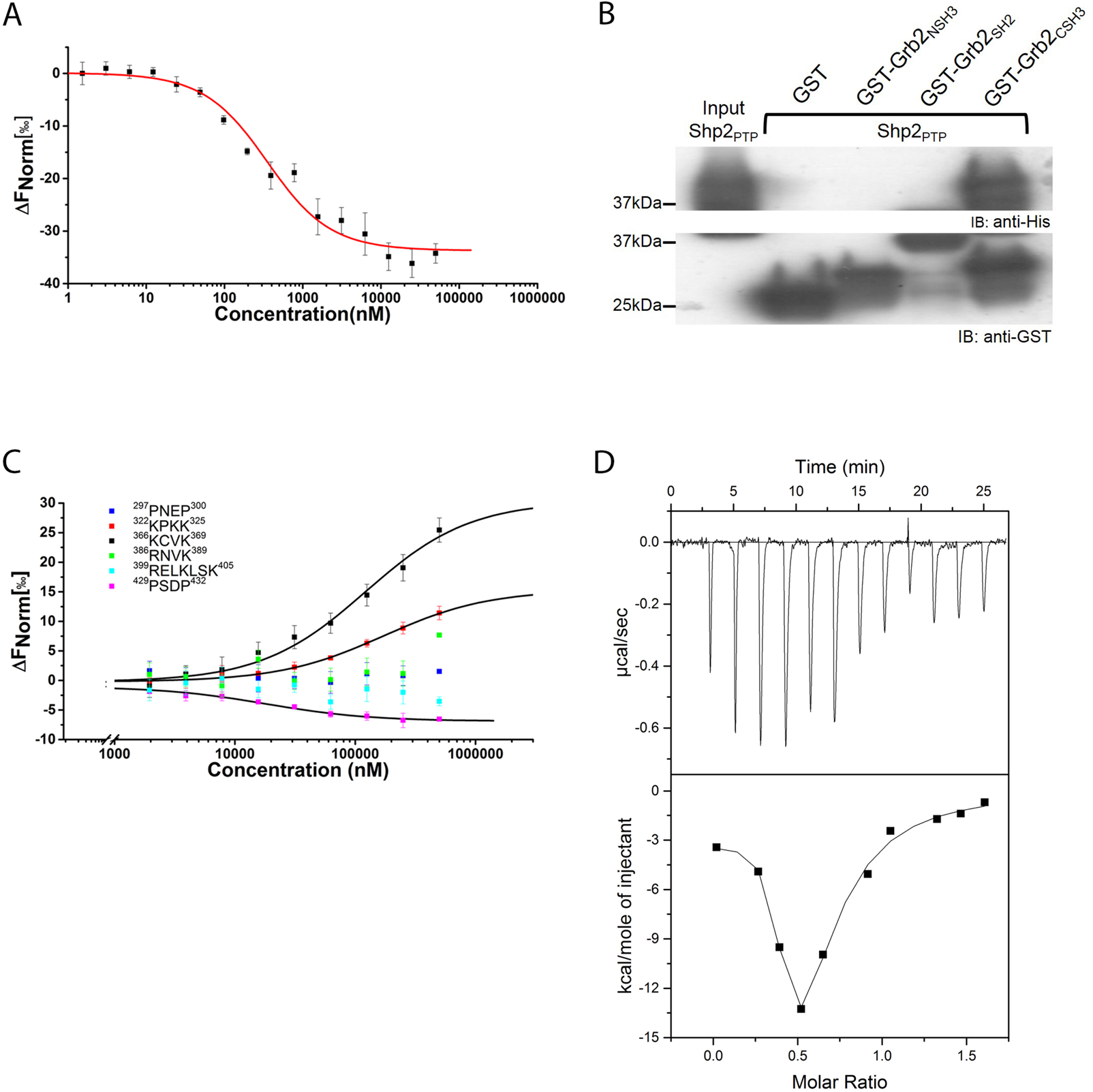
Characterisation of Grb2_Y160E_ binding to Shp2_PTP_ domain. A MST measurement of Shp2_PTP_ binding to fluorescent labelled Grb2_Y160E_ (0.1 µM). Both PTP domain and tandem 2SH2 domains show similar affinity to Grb2_Y160E_. B GST pulldown experiment using GST tagged individual Grb2 domain (NSH3, SH2, and CSH3) to precipitate Shp2 PTP domain. The result clearly indicates the interaction is mediated through Grb2 CSH3 domain. C Sequence analysis identifies 6 potential PxxP or R/KxxK (x is any amino residue) motifs for Grb2 CSH3 domain interaction. Synthesised peptides were used to test the interactions with Grb2 CSH3 domain (Atto 488 labelled, 0.1 µM) using MST. The results show ^322^KPKK^325^, ^366^KCVK^369^, and ^429^PSDP^432^ interact with Shp2 PTP domain. Motif ^429^PSDP^432^ binds to Shp2 PTP domain with a highest affinity of 18 µM. See Table for detailed information and peptide sequences. D ITC was used to corroborate the two binding events between Shp2 and Grb2_Y160E_. Grb2_Y160E_ (200 µM) was titrated into Shp2_Δ69_ (25 µM). Twelve 3 µl injections of Grb2_Y160E_ were titrated into Shp2_Δ69_. Top panel: baseline-corrected power versus time plot for the titration. Bottom panel: the integrated heats and the molar ratio of the Grb2_Y160E_ to Shp2_Δ69_.

Since Grb2 binds to Shp2 through two apparently discrete interactions largely represented by, 1) Grb2_SH2_ binding to Shp2_NSH2_, and 2) Grb2_CSH3_ binding to Shp2_PTP_, we used isothermal titration calorimetry (ITC) to qualitatively corroborate these two binding events. The binding isotherm shows a biphasic profile resulting from an initial binding event with a stoichiometry of 2:1 Shp2_Δ69_:Grb2_Y160E_, which was succeeded by a second binding event with a stoichiometry of unity as more Grb2_Y160E_ was titrated (Fig 3D). These data can be reconciled by a model in which in the initial injections Grb2_Y160E_ is saturated by excess Shp2_Δ69_. Under these conditions Shp2_Δ69_ will occupy both binding sites on Grb2 (i.e. Grb2_SH2_ binding to Shp2_NSH2_ on one Shp2_Δ69_ molecule, and Grb2_CSH3_ binding to Shp2_PTP_ on another Shp2_Δ69_ molecule). On further addition of Grb2_Y160E_, based on the final stoichiometry of 1:1, the complex involves a bidentate interaction in which both the SH2 and CSH3 domain of a single Grb2 concomitantly bind to the NSH2 and PTP domains of a single Shp2 respectively.

### Grb2 increases activity independent of Shp2 phosphorylation

The bidentate binding of Grb2 which includes an interaction with both the NSH2 and the PTP domains of Shp2 implies that complex formation could influence phosphatase activity. From our accumulated data we speculate that the binding of monomeric Grb2 promotes a conformational change releasing Shp2 from the auto-inhibited state. To investigate this, we tagged Shp2_Δ69_ at both the N- and C-termini with blue fluorescent protein (BFP) and green fluorescent protein (GFP) respectively. When this pair of fluorophores are in close proximity (i.e. as would be expected in the auto-inhibited state), excitation of BFP results in fluorescence resonance energy transfer between the two fluorophores, and hence a reduction in the emission intensity from BFP with a concomitant increase from GFP. Conversely, distancing the fluorophores through conformational change reverses this outcome. Steady-state FRET was measured whilst adding an increasing concentration Grb2_Y160E_ to Shp2_Δ69_ (Fig 4A). The data clearly show that, prior to Grb2 addition, the BFP donor emission is low and GFP acceptor emission is high. Recovery of FRET donor BFP emission and a decrease in FRET acceptor GFP emission with increasing concentration of Grb2_Y160E_ results in the growing population of Grb2_Y160E_-Shp2_Δ69_ complex undergoing a conformational change from the auto-inhibited state to a state where the termini of Shp2_Δ69_ are separated.

**Figure 4.**
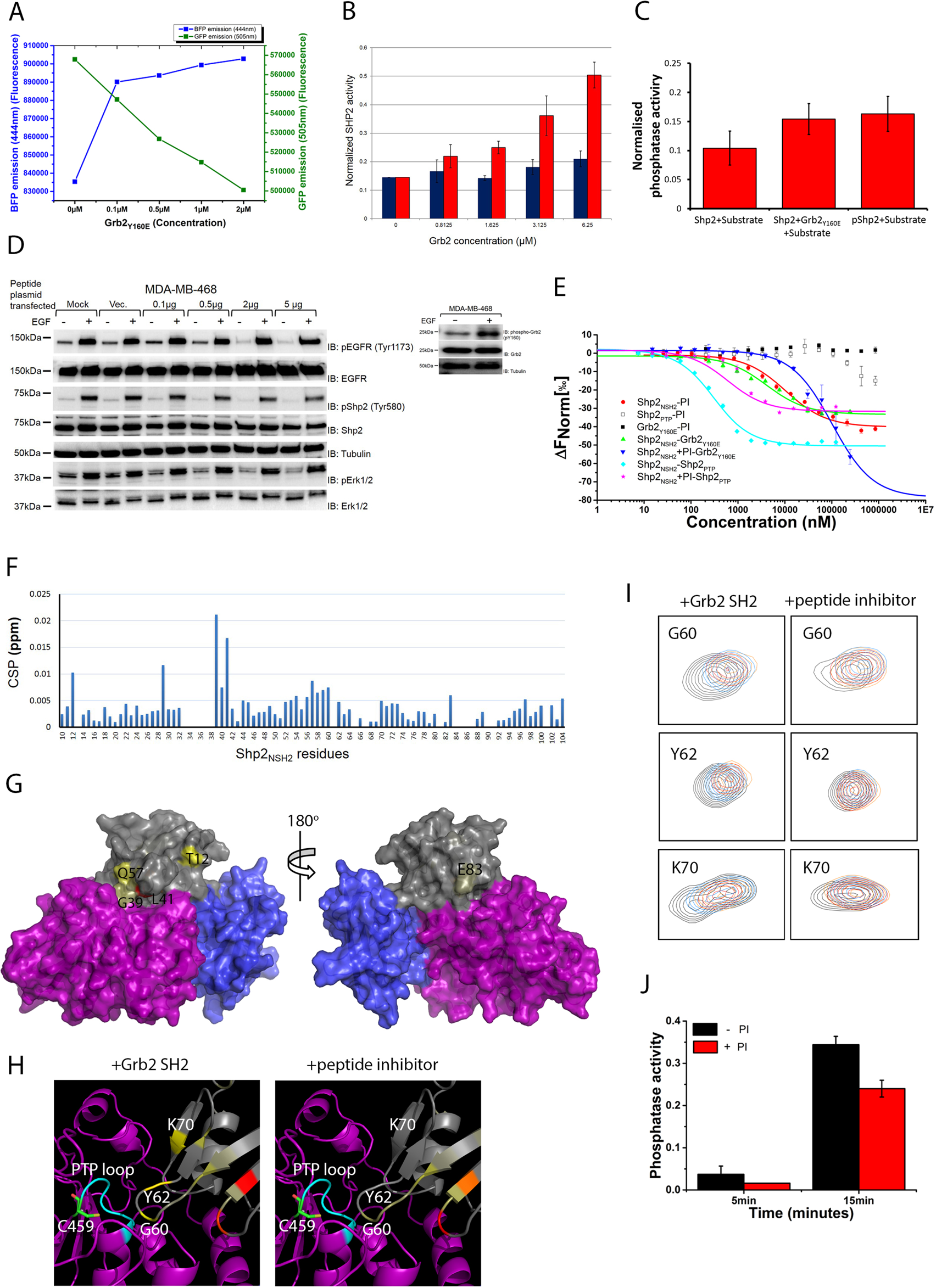
Grb2_Y160E_ upregulates Shp2 activity in a phosphorylation-independent manner. A The effect of mGrb2 binding on Shp2 conformation was studied using steady-state FRET. 0.1µM of Shp2_Δ69_ was N-terminally tagged with a BFP and C-terminally tagged with a GFP. Upon Grb2_Y160E_ titration, the emission of both FRET donor (BFP) and acceptor (GFP) was recorded. B *In vitro* phosphatase assay using recombinant-unphosphorylated Shp2_Δ69_ (0.1 µM) and Grb2 (WT or Y160E). Free phosphate generated from hydrolysis of the pY from the substrate peptide (ENDpYINASL) was measured by the absorbance of a malachite green molybdate phosphate complex. The increasing concentration of Grb2_Y160E_ gradually enhances Shp2 activity while the dimeric Grb2 has no effect on Shp2 activity. C *In vitro* phosphatase assay using both recombinant-unphosphorylated (0.1 µM) and phosphorylated (0.1 µM) full length Shp2 to compare the degree of enhanced phosphatase activity mediated by mGrb2 (10 µM) binding. This assay demonstrates that the mGrb2 binding –induced enhancement of Shp2 activity is comparable to the phosphorylated Shp2. D Overexpression of Peptide Inhibitor (PI) in MDA-MD-468 cells decreases Shp2-mediated MAPK signalling in the absence of EGF ligand stimulation. E Biophysical characterisation of the peptide inhibitor (PI) using MST. Red: Atto 488 labelled Shp2_NSH2_ (0.1 µM) binds to PI. Empty Square: Atto 488 labelled Shp2_PTP_ (0.1 µM) does not bind to PI. Black: Atto 488 labelled Grb2_Y160E_ (0.1 µM) does not bind to PI. Green: Atto 488 labelled Shp2_NSH2_ (0.1 µM) binds to Grb2_Y160E_. Blue: Atto 488 labelled Shp2_NSH2_ (0.1 µM) was saturated with 10 µM of PI. PI-bound Shp2_NSH2_ (0.1 µM) decrease the binding to Grb2_Y160E_. Cyan: Atto 488 labelled Shp2_NSH2_ (0.1 µM) binds to Shp2_PTP_. Magenta: binding of PI does not affect labelled Shp2_NSH2_ (0.1 µM() binding to Shp2_PTP_. F The NMR HSQC titration results indicate that the predominant CSPs on Shp2_NSH2_ occur on those residues involved in the auto-inhibition interface. This strongly suggest that binding of mGrb2 results in the conformational change of Shp2 and releases it form it auto-inhibition state (left panel). However, binding of Peptide Inhibitor (PI) results in the less CSPs within the auto-inhibition interface (right panel). G Mapping on the surface of Shp2 (PDB: 2SHP) of the consensus residues (painted light yellow to red) exhibiting strong CSPs upon PI binding. The Shp2_NSH2_ domain is in grey, Shp2_CSH2_ domain is in blue, and the SHp2_PTP_ domain is in purple. This identify the potential binding sites on Shp2_NSH2_ for PI binding. H Comparison of residues G60, Y62, and K70 chemical shifts in the presence of Grb2_SH2_ (left panel) or PI (right panel). Overlay of a region of the ^15^N-^1^H HSQC (heteronuclear single-quantum coherence) spectra of Shp2_NSH2_ alone (black) and of Shp2_NSH2_ and its complex (red). Binding of PI induces less CSPs on residues G60, Y62, and K70. I Interaction of ^15^N-labelled Shp2_NSH2_ and peptide inhibitor (PI) determined by NMR chemical shift perturbation (CSP) plot for the interaction of PI to ^15^N-labelled Shp2_NSH2_. Minimal chemical shift perturbation upon PI mapped on Shp2_NSH2_ sequence. J Using an *in vitro* phosphatase assay it is demonstrated that the PI inhibits mGrb2-induced upregulation of Shp2 (0.1 µM) phosphatase activity. Black: Shp2 activity towards a substrate peptide in the presence of mGrb2 (10 µM). Red: Shp2 activity towards a substrate peptide in the presence of mGrb2 (10 µM) and PI (10µM).

To assess whether the Grb2 binding-induced conformational change affected enzymatic activity we conducted an *in vitro* phosphatase assay. A pY-containing peptide substrate (ENDpYINASL) was incubated with non-phosphorylated Shp2_Δ69_ and different concentrations of Grb2_Y160E_. Free phosphate generated from hydrolysis of the pY was measured by the absorbance of a malachite green molybdate phosphate complex (Fig 4B). A significant increase in phosphatase activity was measured with the increasing presence of Grb2_Y160E_. Importantly, a similar level of turnover of the peptide was observed on comparing the monomeric Grb2-induced phosphatase activity to that observed for phosphorylated Shp2 in the absence of the adaptor protein (pShp2; Fig 4C). These *in vitro* experiments suggest that under basal conditions Shp2 activity can be up-regulated through binding to monomeric Grb2 alone, and that this activity is at least as high as occurs when the phosphatase is phosphorylated as seen in growth factor-stimulated cells.

### Monomeric Grb2 is associated with the up-regulation of Shp2 in cancer

Shp2 activity has been shown to play a key role in cancer progression. Since mGrb2 can promote phosphatase activity in the absence of Shp2 phosphorylation, this could lead to a proliferative outcome in cells through up-regulation of the Erk1/2 pathway without the need for the elevated kinase activity often associated with cancer phenotypes. The role of up-regulation of non-phosphorylated Shp2 through binding to mGrb2 in cancer has not been previously been investigated. For example, in the triple negative breast cancer cell line MDA-MB-468 in the absence of extracellular stimulation we see negligible background pShp2. However, there is a significant concentration of pErk1/2 (Fig 4D, compare lane 1 and lane 2). These cells have a background level of monomeric phosphorylated Grb2. Indeed, Grb2 phosphorylation under basal non-stimulatory conditions has been reported for a number of cell lines (Ahmed et al., 2013; Fig 4D inset), suggesting that a prevailing mGrb2-Shp2 interaction could permit Shp2-mediated signalling.

To ascertain whether the elevated pErk1/2 was accrued from increased phosphatase activity resulting from the interaction of Grb2 with Shp2 we identified a polypeptide inhibitor (PI; Appendix S2A), which bound to Shp2 and abrogated Grb2 binding. PI binds to Shp2_NSH2_ (K_d_ = 7.24 ± 0.52 µM) and was shown not to bind to either full-length Grb2 or Shp2_PTP_ (Fig 4E and Table). Pre-incubating Shp2_NSH2_ with PI inhibits the mGrb2-Shp2_NSH2_ interaction reducing the affinity by approximately 50-fold (K_d_ = 101 ± 6 µM; compared to K_d_ = 2.3 ± 0.2 µM; Fig 4E). The inhibitory effect of PI keeps Shp2 in its auto-regulated state because its presence does not impact on the binding of Shp2_PTP_ to Shp2_NSH2_ in the closed state (Shp2_PTP_ + Shp2_NSH2_, K_d_ = 0.25 µM; Shp2_PTP_ + Shp2_NSH2_ + PI, K_d_ = 0.55 µM; Fig 4E and Table). NMR CSP mapping revealed that the binding interface between Shp2_NSH2_ and PI overlapped with that of Grb2_SH2_ (Fig 4F and 4G). Importantly, the binding of PI does not seem to affect the residues in the interface of the intramolecular NSH2-PTP domain interaction, hence Shp2 is able to maintain its closed, auto-inhibited state (Fig 4H and 4I). The ability of PI to inhibit up-regulation of Shp2 by blocking the interaction with Grb2, but maintaining the intramolecular auto-inhibition interaction is exemplified by showing its ability to reduce phosphatase activity (Fig 4J). Therefore, the PI provides a useful ‘bio-tool’ to support the mechanistic detail, as well as enabling us to probe the effect of inhibiting activation of Shp2 by Grb2.

Using our novel PI bio-tool we can demonstrate that inhibition of the mGrb2-Shp2 interaction through increased doses of PI, resulted in a reduction in Erk1/2 signalling in MDA-MB-468 breast cancer cells (Fig 4D). Stimulation of the cells with EGF negates the effect of inhibition of the mGrb2 activation of Shp2. This results from up-regulation of kinase activity in the cells as is clearly seen by the appearance of pShp2 and the significant appearance pErk1/2 even in the presence of PI. After 15 minutes in the phosphatase assay the activity is reduced by approximately one third in the presence of the PI. These data provide compelling evidence for the ability of the Grb2-Shp2 driven up-regulation of Erk1/2 signalling to contribute to the cancer pathology represented by this cell line.

## Discussion

Kinase and phosphatase activity has to be precisely controlled to limit aberrant signal transduction in cells. It has previously been demonstrated that Shp2 function is dependent on RTK activity, and is accompanied by phosphorylation of tyrosine residues Y542 and Y580 which appears to facilitate the conformational change that abrogates an intramolecular interface between the NSH2 and PTP domains of Shp2 and relieves the auto-inhibited state. Activated Shp2 can drive proliferative signalling through the Erk1/2 pathway. In this work we show that the up-regulation of Shp2 can also be accomplished in the absence of RTK activity, through the binding of the adaptor protein Grb2. The SH2 and CSH3 domains of Grb2 form a bidentate interaction with the NSH2 and PTP domains of Shp2 respectively. Interaction with Shp2 in this way results in a Grb2-dependent conformational change which opens the auto-inhibited state to facilitate phosphatase activity.

The relevance of a mechanism for Shp2 up-regulation in non-stimulated cells is likely to be associated with the requirement of cells to maintain homeostatic and metabolic ‘house-keeping’ function in the absence of extracellular stimulation. We have previously shown that similar functional activation of enzymes and adaptors associated with RTK-mediated signalling pathways occurs under basal conditions without the need for kinase up-regulation (e.g. Plcγ1, Timsah et al., 2014; Shc, Suen et al., 2014). This second tier of signalling, below the profound and defined effects of full kinase activation resulting from extracellular stimulation, is likely to be important in maintaining cell viability and in responding to stress. Aberrancies in second tier signalling can also be associated with cancer pathology (Timsah et al 2014). This appears to be reflected here where, through interaction with mGrb2, Shp2 can be engaged in signalling in cancer cells without the need for mutated, dysfunctional kinases. Therefore, in cells depleted for a given RTK, as is found in triple-negative breast cancer (e.g. in MDA-MB-468 cells which do not express Her2; Fig 4D), proliferative or metastatic signalling could be driven by uncontrolled impact of mGrb2-mediated Shp2 activation. The importance of the interaction with Grb2 may also be emphasised by the observation of two reported oncogenic Shp2 mutations in the NSH2 which are localised to the identified Grb2 binding site, namely T42A and N58K. Change-of-function associated with these mutations would potentially release regulatory control of background tyrosine kinase-mediated signalling.

Non-stimulatory conditions portray many features of the early phases of tumour development and progression as well as prevailing in resistant cells treated with kinase inhibitors. Thus, the understanding of how signalling proteins behave and are controlled under these conditions is the necessary foundation to develop more efficient therapeutic strategies. It remains to be seen whether examples of mGrb2-mediated activation of Shp2 are common regulators of signalling in basal cells, and particularly whether aberrancies resulting from environmental change-related stress can drive tumorigenesis through perturbation of Grb2 concentration. Nonetheless, we have shown that inhibition of the mGrb2-Shp2 interaction can down-regulate Shp2 activity, and help to sustain the auto-inhibited form of the phosphatase. The binding of a polypeptide to the NSH2 domain reduced activity *in vitro* and reduced pErk1/2 in MDA-MB-468 cells. Thus, for down-regulation of aberrant signalling in non-stimulated cells, the development of therapeutics based on the PI could be an important novel route to intervention.

## Materials and Methods

### Cell culture

HEK293T was maintained in DMEM (Dulbecco’s modified Eagle’s high glucose medium) supplemented with 10% (v/v) FBS (foetal bovine serum) and 1% antibiotic/antimycotic (Lonza) in a humidified incubator with 10% CO_2_.

### Plasmids

For recombinant protein production in *E coli*, genes of interest were amplified using standard PCR methods. Following designated restriction enzyme digestions, fragments were ligated into pET28b or pGEX4T1. For cell-based strep-tag pulldown experiments, Grb2 genes (wild type or Y160E) were amplified with an N-terminal strep-tag and cloned in a pcDNA6 vector. For the cellular FLIM study, Grb2 genes were cloned in pECFP, and pcDNA-RFP vectors.

### Protein expression and purification

Proteins were expressed in BL21 (DE3) cells. 20 ml of cells grown overnight were used to inoculate 1 litre of LB media with antibiotic (50 µg/ml kanamycin for pET28b-backboned plasmids or 50 µg/ml ampicillin for pGEX4T1-backboned plasmids). The culture was grown at 37 °C with constant shaking (200 rpm) until the OD_600_ = 0.7. At this point the culture was cooled down to 18 °C and 0.5 mM of IPTG was added to induce protein expression for 12 h before harvesting. Harvested cells were suspended in Talon buffer A (20mM Tris, 150mM NaCl, and 1mM β-ME, at pH 8.0) and lysed by sonication. Cell debris was removed by centrifugation (20,000 rpm at 4°C for 1 h). The soluble fraction was applied to an Akta Purifer System for protein purification. Elution was performed using Talon buffer B (20mM Tris, 150mM NaCl, 200mM imidazole and 1mM β-ME at pH 8.0). Proteins were concentrated to 2 ml and applied to a Superdex SD75 column using a HEPES buffer at pH 7.5 (20 mM HEPES, 150mM NaCl, and 1mM TCEP, at pH 7.5).

### Pulldown/Precipitation

Purified protein or total HEK293T cell lysate were prepared in 1 ml volume for GST pulldown. 50 μl of 50% glutathione beads were added and incubated for overnight. The beads were spun down at 5,000 rpm for 3 minutes, supernatant was removed and the beads were washed with 1 ml lysis buffer. This washing procedure was repeated five times in order to remove non-specific binding. After the last wash, 50 μl of 2x Laemmli sample buffer were added, the samples were boiled and subjected to SDS-PAGE and western blot assay.

### Fluorescence Lifetime Imaging Microscopy, FLIM

HEK293T cells were co-transfected with cyan fluorescent protein- and red fluorescent protein-tagged Grb2 and Shp2 respectively (CFP-Grb2 and RFP-Shp2). After 24 h cells were seeded onto glass coverslips and allowed to grow for a further 24 h, fixed by the addition of 4% (w/vol) paraformaldehyde (PFA) pH 8.0. Following 20 min incubation at room temperature cells were washed 6-7 times with PBS, pH 8.0. Coverslips were mounted onto a slide with mounting medium (0.1% p-phenylenediamine and 75% glycerol in PBS at pH 7.5–8.0). FLIM images were captured using a Leica SP5 II confocal microscope with an internal FLIM detector. CFP was excited at 860 nm with titanium–sapphire pumped laser (Mai Tai BB, Spectral Physics) with 710-920 nm tunability and 70 femtosecond pulse width. A Becker & Hickl SPC830 data and image acquisition card was used for time-correlated single-photon counting (TCSPC); electrical time resolution was 8 picoseconds with a pixel resolution of 256 x 256. Data processing and analysis were performed using a B&H SPC FLIM analysis software. The fluorescence decays were fitted to a single exponential decay model.

### Isothermal Titration Calorimetry, ITC

ITC experiments were carried out using a MicroCal iTC200 instrument (Malvern), and data were analysed using ORIGIN7 software. To avoid heats associated with protein dissociation Grb2_Y160E_ was titrated into Shp2_Δ69_ at 25°C. The heat per injection was determined and subtracted from the binding data. Data was analysed using a single independent site model using the Origin software.

### Microscale Thermophoresis, MST

The Grb2 and Shp2 interactions were measured using the Monolith NT.115 MST instrument from Nanotemper Technologies. Proteins were fluorescently labelled with Atto 488 NHS ester (Sigma) according to the manufacturer’s protocol. Labelling efficiency was determined to be 1:1 (protein to dye) by measuring the absorbance at 280 nm and 488 nm. A solution of unlabelled protein was serially diluted in the presence of 100 nM labelled protein. The samples were loaded into capillaries (Nanotemper Technologies). Measurements were performed at 25°C in 20 mM HEPES buffer, pH 7.5, with 150 mM NaCl, 0.5 mM TCEP, and 0.005% Tween 20, Data analyses were performed using Nanotemper Analysis software, v.1.5.41 and plotted using OriginPro 9.1.

### Biolayer Interferometry, BLI

BLI experiments were performed using a FortéBio Octet Red 384 using Anti-GST sensors. Assays were done in 384 well plates at 25 °C. Association was measured by dipping sensors into solutions of analyte protein for 120 seconds and was followed by moving sensors to wash buffer for 120 seconds to monitor the dissociation process. Raw data shows a rise in signal associated with binding followed by a diminished signal after application of wash buffer.

### *In Vitro* Phosphatase Assay

*In vitro* Shp2 activity assays were carried out using the Tyrosine Phosphatase Assay System (Promega) according to the manufacturer’s manual. In brief, recombinant Shp2_Δ69_ was mixed with the phospho-peptide substrate in the presence or absence of different concentrations of Grb2 (Grb2_WT_ or Grb2_Y160E_). The method is based on measuring the absorbent change generated after formation of a reaction mixture of molybdate:malachite green-phosphate complex of the free phosphate.

### Nuclear Magnetic Resonance Spectroscopy, NMR

Backbone resonances for the ^1^H, ^15^N, ^13^C labelled sample of Shp2 were assigned by recording a standard Bruker or BEST TROSY version (Lescop et al., 2007) of 3D backbone resonance assignment spectra (HNCA, HNCOCA, HNCACB, CACBCONH, HNCO and HNCACO) with NUS sampling technique (25% fraction). The titration experiments of Grb2 and PI into Shp2 were recorded using ^15^N HSQC pulse sequence from Bruker standard library. For that, samples of 200 µM uniformly ^15^N-labelled Shp2 in20mM HEPES, 150mM NaCl, 0.5mM TCEP. pH7.5 and 10% (v/v) D2O with 1:1, 1:2, 1:3, 1:4, 1:5, 1:6 molar ratio of Shp2:Grb2 and 1:1, 1:2, 1:5, 1:10, 1:20 molar ratio of Shp2:IP1 were prepared. All measurements were recorded at 25°C using a Brüker Avance III 750 MHz and 950 MHz spectrometers equipped with a Bruker TCI triple-resonance cryogenically coiled probes. Data was processed with NMRPipe (Delaglio et al., 1995) and analyzed with CcpNmr Analysis software package (Vranken et al., 2005). Chemical shift perturbations (CSPs) for individual residues were calculated from the chemical shift for the backbone amide ^1^H (Δω_H_) and ^15^N (Δω_N_) using the following equation: 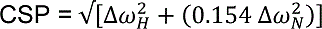
 (Evenas et al., 2001).

## ACKNOWLEDGEMENTS

This work was funded by Cancer Research UK (Grant C57233/A22356). We thank Steve Homans for his assistance on interpretation of NMR data and Amy Stainthorp (University of Leeds, School of Molecular and Cellular Biology) for critical discussion and useful comments.

## AUTHOR CONTRIBUTIONS

C-CL and JEL conceived of the project and designed experiments. C-CL, KMS, and ZA carried out experiments. LW, and AK analysed NMR data. C-CL and JEL wrote the paper with input from all authors.

---

**Figure EV1:**
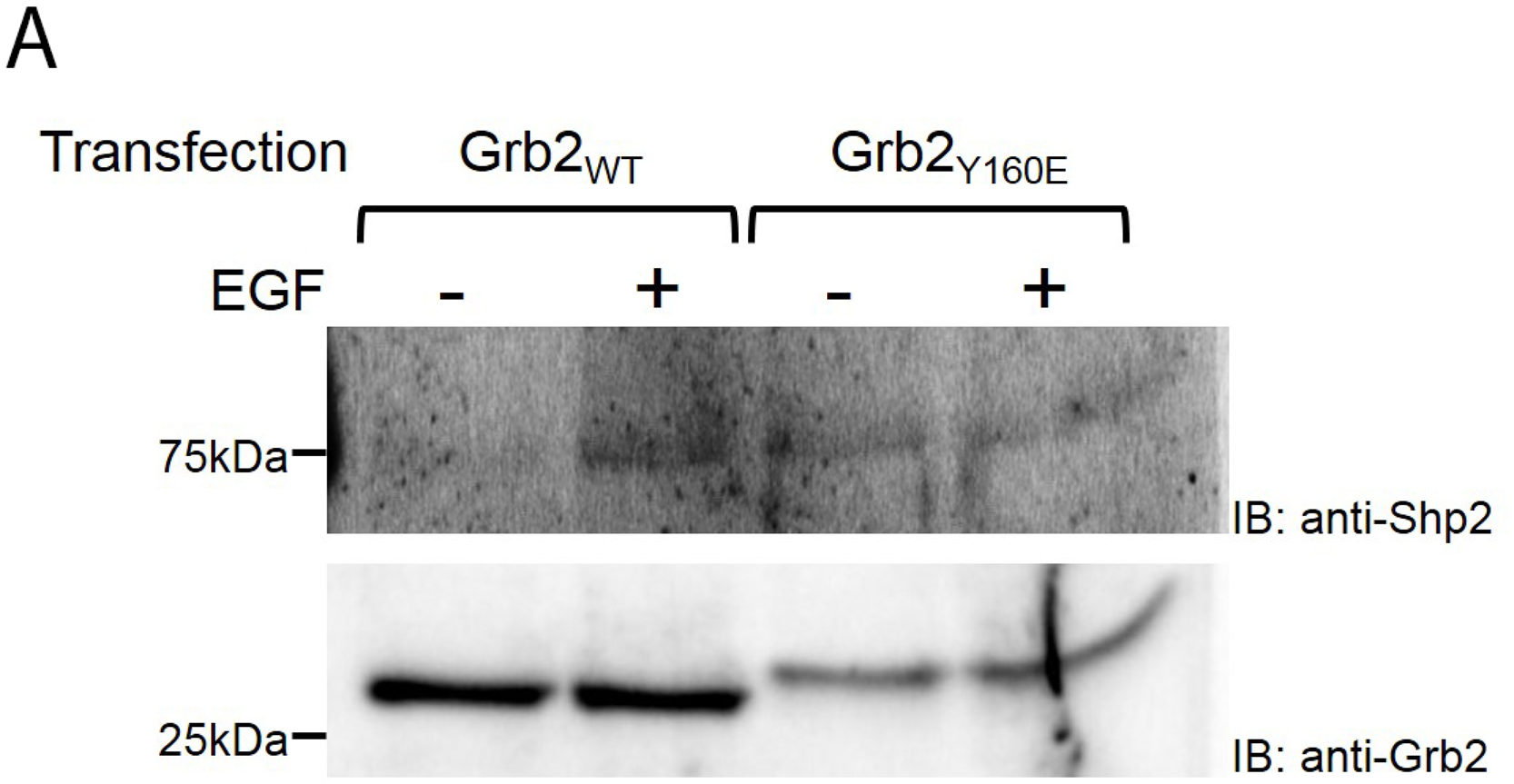
Interaction of Shp2 with monomeric Grb2 at basal state. (A) Strep-tagged Grb2 plasmids (dimeric Grb2, Grb2_WT_; monomeric Grb2, Grb2_Y160E_) were transfected into HEK293T cells and cells were remained serum-starved or EGF-stimulated. Grb2-Shp2 complexes were pulldown using streptavidin beads and precipitated Shp2 was immunoblotted using a Shp2 antibody. Both dimeric and monomeric Grb2 can precipitate Shp2 upon EGF stimulation. However, in the absence of EGF stimulation, only monomeric Grb2 (Grb2_Y160E_) can precipitate Shp2.

**Figure EV2:**
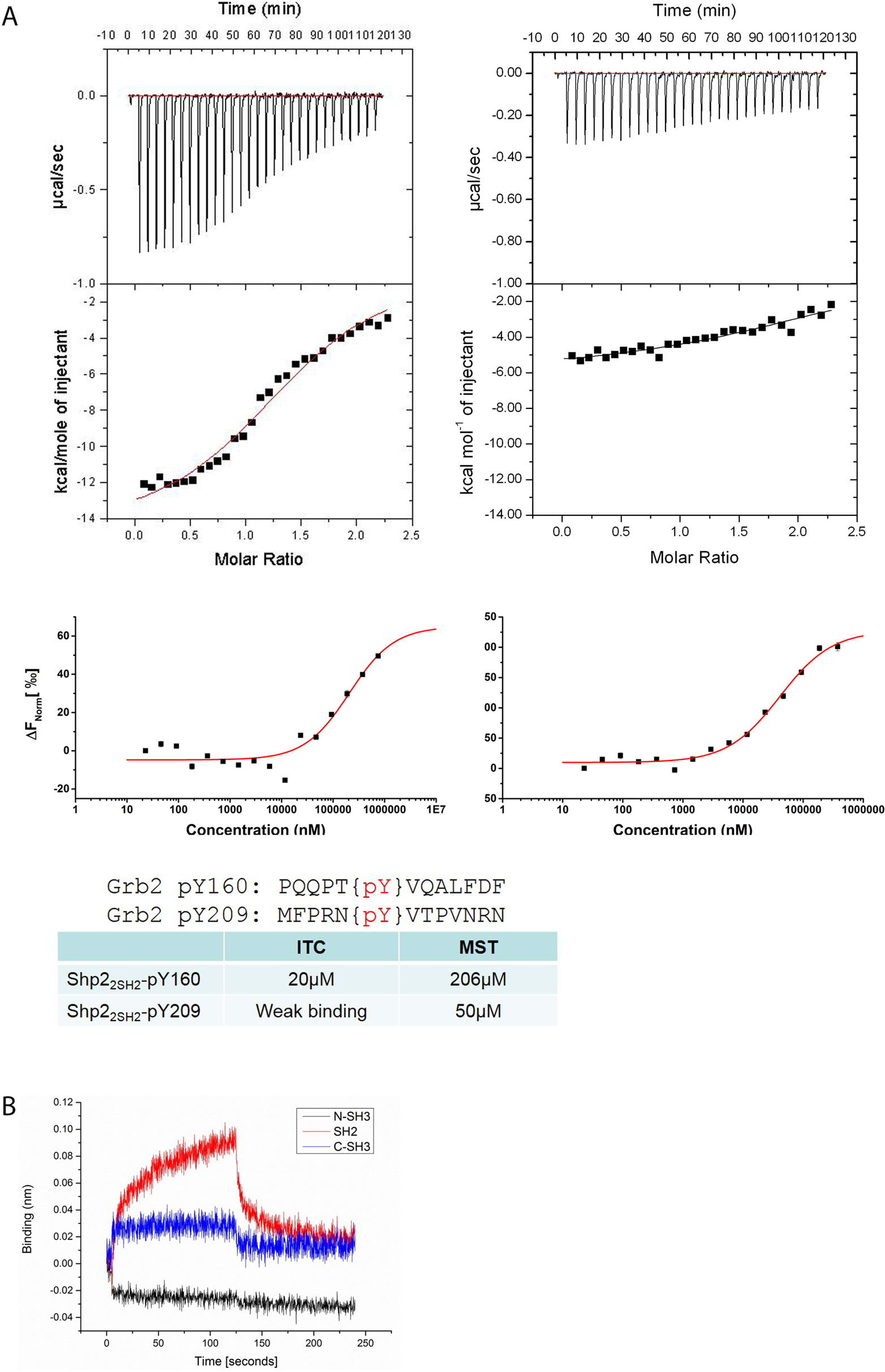
Phosphorylation-dependent interaction is negligible in Shp2_2SH2_ and mGrb2 interaction. (A) Phosphorylation of Grb2 Y160 results in the dimer dissociation. Although there is no trace of phosphorylation on the phosphorylation mimic Grb2_Y160E_, we performed experiments to confirm the interaction is phosphorylation-independent. Y160 and Y209 are two major phosphorylation sites on Grb2; synthesised phospho-peptides that bear PY160 or PY209 were used for biophysical characterisation. The PY 160 peptide, shows an affinity of 20 µM in ITC measurement and 206 µM in MST measurement. Bot determined affinity are at least 70-fold weaker than the phosphorylation-independent Shp2_2SH2_ and Grb2_Y160E_ interaction (0.28 ± 0.03 µM). Similar to the PY160 peptide, PY209 peptide has a 180-fold weaker affinity comparing to Shp2_2SH2_ and Grb2_Y160E_ interaction in the MST measurement. (B) In order to determine the domain in Grb2 that is responsible for Shp2_NSH2_ binding, GST-tagged Shp2_NSH2_ was immobilised on a BLI sensor and individual Grb2 domains (Black: NSH3, Red: SH2, and Blue: CSH3) were used to characterise the interaction. The BLI screen results clearly indicates that the SH2 domain of Grb2 is required for binding to Shp2_NSH2_.

**Figure EV3:**
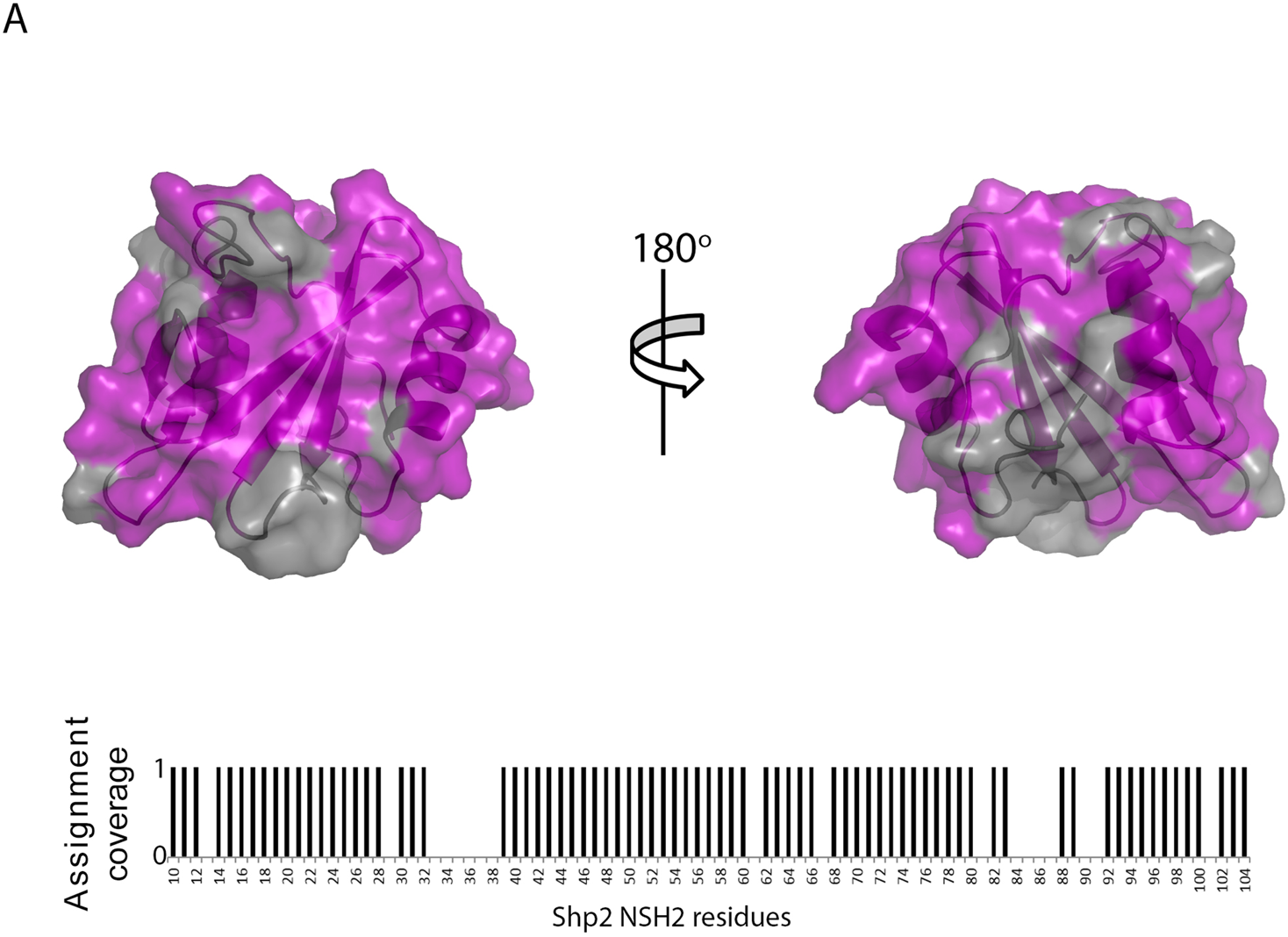

## REFERENCES

1. Aceto N, Sausgruber N, Brinkhaus H, Gaidatzis D, Martiny-Baron G, Mazzarol G, Confalonieri S, Quarto M, Hu G, Balwierz PJ, Pachkov M, Elledge SJ, van Nimwegen E, Stadler MB, Bentires-Alj M (2012) Tyrosine phosphatase SHP2 promotes breast cancer progression and maintains tumor-initiating cells via activation of key transcription factors and a positive feedback signaling loop. Nat Med 18: 529–537

2. Ahmed Z, George R, Lin C-C, Suen KM, Levitt JA, Suhling K, Ladbury JE (2010) Direct binding of Grb2 SH3 domain to FGFR2 regulates SHP2 function. Cell Signal 22: 23–33

3. Ahmed Z, Lin C-C, Suen KM, Melo FA, Levitt JA, Suhling K, Ladbury JE (2013) Grb2 controls phosphorylation of FGFR2 by inhibiting receptor kinase and Shp2 phosphatase activity. J Cell Biol 200: 493–504

4. Ahmed Z, Timsah Z, Suen KM, Cook NP, Lee IV GR, Lin C-C, Gagea M, Marti AA, Ladbury JE (2015) Grb2 monomer–dimer equilibrium determines normal versus oncogenic function. Nat Commun 6: 8007

5. Araki T, Nawa H, Neel BG (2003) Tyrosyl phosphorylation of Shp2 is required for normal ERK activation in response to some, but not all, growth factors. J Biol Chem 278: 41677–41684

6. Bentires-Alj M, Gil SG, Chan R, Wang ZC, Wang Y, Imanaka N, Harris LN, Richardson A, Neel BG, Gu H (2006) A role for the scaffolding adapter GAB2 in breast cancer. Nat Med 12: 114–121

7. Bentires-Alj M, Paez JG, David FS, Keilhack H, Halmos B, Naoki K, Maris JM, Richardson A, Bardelli A, Sugarbaker DJ, Richards WG, Du J, Girard L, Minna JD, Loh ML, Fisher DE, Velculescu VE, Vogelstein B, Meyerson M, Sellers WR, Neel BG (2004) Activating mutations of the noonan syndrome-associated SHP2/PTPN11 gene in human solid tumors and adult acute myelogenous leukemia. Cancer Res 64: 8816–8820

8. Chan RJ, Feng GS (2007) PTPN11 is the first identified proto-oncogene that encodes a tyrosine phosphatase. Blood 109: 862–867

9. Chan G, Kalaitzidis D, Neel BG (2008) The tyrosine phosphatase Shp2 (PTPN11) in cancer. Cancer Metastasis Rev 27: 179–192

10. Cunnick JM, Mei L, Doupnik CA, Wu J (2001) Phosphotyrosines 627 and 659 of Gab1 constitute a bisphosphoryl tyrosine-based activation motif (BTAM) conferring binding and activation of SHP2. J Biol Chem 276: 24380–24387

11. Delaglio F, Grzesiek S, Vuister GW, Zhu G, Pfeifer J, Bax A. NMRPipe: a multidimensional spectral processing system based on UNIX pipes. (1995) J. Biomol. NMR 6: 277–293.

12. Eck MJ, Pluskey S, Trub T, Harrison SC, Shoelson SE (1996) Spatial constraints on the recognition of phosphoproteins by the tandem SH2 domains of the phosphatase SH-PTP2. Nature 379: 277–280

13. Evenäs J, Tugarinov V, Skrynnikov NR, Goto NK, Muhandiram R, Kay LE. (2001). Ligand-induced structural changes to maltodextrin-binding protein as studied by solution NMR spectroscopy. J. Mol. Biol. 309: 961–974

14. Feng GS (1999) Shp-2 tyrosine phosphatase: signaling one cell or many. Exp Cell Res 253: 47–54

15. Hof P, Pluskey S, Dhe-Paganon S, Eck MJ, Shoelson SE (1998) Crystal structure of the tyrosine phosphatase SHP-2. Cell 92: 441–450

16. Keilhack H, David FS, McGregor M, Cantley LC, Neel BG (2005) Diverse biochemical properties of Shp2 mutants. Implications for disease phenotypes. J Biol Chem 280: 30984–30993

17. Lechleider RJ, Sugimoto S, Bennett AM, Kashishian AS, Cooper JA, Shoelson SE, Walsh CT, Neel BG (1993) Activation of the SH2-containing phosphotyrosine phosphatase SH-PTP2 by its binding site, phosphotyrosine 1009, on the human platelet-derived growth factor receptor. J Biol Chem 268: 21478–21481

18. Lescop E, Schanda P, Brutscher B. (2007). A set of BEST triple-resonance experiments for time-optimized protein resonance assignment. J Magn Reson 187: 163–169.

19. Lu W, Gong D, Bar-Sagi D, Cole PA (2001). Site-specific incorporation of a phosphotyrosine mimetic reveals a role for tyrosine phosphorylation of SHP-2 in cell signaling. Mol Cell 8: 759–769

20. Lin C-C, Melo FA, Ghosh R, Suen KM, Stagg LJ, Kirkpatrick J, Arold ST, Ahmed Z, Ladbury JE (2012) Inhibition of basal FGF receptor signaling by dimeric Grb2. Cell 149: 1514–1524

21. Neel BG, Tonks NK (1997) Protein tyrosine phosphatases in signal transduction. Curr Opin Cell Biol 9: 193–204

22. O’Reilly AM, Pluskey S, Shoelson SE, Neel BG (2000) Activated mutants of SHP-2 preferentially induce elongation of Xenopus animal caps. Mol Cell Biol 20: 299–311

23. Pluskey S, Wandless TJ, Walsh, CT, Shoelson SE (1995) Potent stimulation of SH-PTP2 phosphatase activity by simultaneous occupancy of both SH2 domains. J Biol Chem 270: 2897–2900

24. Sun J, Lu S, Ouyang M, Lin LJ, Zhuo Y, Liu B, Chien S, Neel BG, Wang Y (2013) Antagonism between binding site affinity and conformational dynamics tunes alternative cis-interactions within Shp2. Nat Commun 4: 2037

25. Sugimoto S, Wandless TJ, Shoelson SE, Neel BG, Walsh, CT (1994) Activation of the SH2-containing protein tyrosine phosphatase, SH-PTP2, by phosphotyrosine-containing peptides derived from insulin receptor substrate-1. J Biol Chem 269: 13614–13622

26. Timsah Z, Ahmed Z, Lin C–C, Melo FA, Stagg LJ, Leonard PG, Jeyabal P, Berrout J, O’Neil RG, Bogdanov M, Ladbury J.E. (2014) Competition between Grb2 and Plcγ1 for binding to FGFR2 regulates constitutive phospholipase activity and invasive response. Nature Struct Mol Biol 21: 180–188.

27. Tonks NK, Neel BG (2001) Combinatorial control of the specificity of protein tyrosine phosphatases. Curr Opin Cell Biol 13: 182–195

28. Van Vactor D, O’Reilly AM, Neel BG (1998) Protein tyrosine phosphatases in the developing nervous system. Curr Opin Genet Dev 8: 112–126

29. Vranken WF, Boucher W, Stevens TJ, Fogh RH, Pajon A, Llinas M, Ulrich EL, Markley JL, Ionides J, Laue ED (2005). The CCPN data model for NMR spectroscopy: development of a software pipeline. *Proteins: Struct, Funct*, & Bioinform. 59: 687–696

